# The Exon Junction Complex and intron removal prevents resplicing of mRNA

**DOI:** 10.1101/2020.07.31.231498

**Authors:** Brian Joseph, Eric C. Lai

## Abstract

Accurate splice site selection is critical for fruitful gene expression. Here, we demonstrate the *Drosophila* EJC suppresses hundreds of functional cryptic splice sites (SS), even though majority of these bear weak splicing motifs and appear incompetent. Mechanistically, the EJC directly conceals splicing elements through position-specific recruitment, preventing SS definition. We note that intron removal using strong, canonical SS yields AG|GU signatures at exon-exon junctions. Unexpectedly, we discover that scores of these minimal exon junction sequences are in fact EJC-suppressed 5’ and 3’ recursive SS, and that loss of EJC regulation from such transcripts triggers faulty mRNA resplicing. An important corollary is that intronless cDNA expression constructs from aforementioned targets yield high levels of unanticipated, truncated transcripts generated by resplicing. Consequently, we conclude the EJC has ancestral roles to defend transcriptome fidelity by (1) repressing illegitimate splice sites on pre-mRNAs, and (2) preventing inadvertent activation of such sites on spliced segments.

## Introduction

The generation of mature mRNA involves multiple metabolic events, which alter the structure of the messenger ribonucleoprotein particle (mRNP). Since the late 1980s, there has been growing appreciation that one such event, splicing, both modulates mRNP composition and influences downstream metabolic processes. Notably, the Exon Junction Complex (EJC) has emerged as a key, splicing-dependent hub for both mRNA processing and post-transcriptional gene regulation.

The EJC is a multisubunit conglomerate that is deposited in a sequence-independent fashion ∼24 nt upstream of exon-exon junctions (Boehm and Gehring 2016; Le Hir et al. 2016). Assembly of its three-member core complex begins during splicing, and the first step involves the position-specific deposition of the DEAD-box protein eIF4AIII onto RNA by the spliceosome factor CWC22. Next, a heterodimer of MAGOH/Mago Nashi and RBM8A/Y14/Tsunagi binds eIF4AIII, stabilizing the complex on RNA. The core EJC complex interacts with multiple peripheral complexes involved in diverse RNA metabolism pathways. These include CASC3/Barentsz, members of the ASAP/PSAP (ACIN1/PNN-RNPS1-SAP18) complexes and other splicing factors, the export factor ALYREF, and the NMD protein UPF3B (Schlautmann and Gehring 2020). Accordingly, EJC dysfunction broadly affects development, disease and cancer (Bonnal et al. 2020).

Curiously, while the EJC is well-conserved, the literature indicates fundamental differences in its requirements between invertebrates and vertebrates (Schlautmann and Gehring 2020). The EJC was first linked to the process of nonsense mediated mRNA decay (Kim et al. 2001; Lykke-Andersen et al. 2001), a process that exploits deposition of the EJC by the spliceosome (Le Hir et al. 2000). Translation removes EJCs from the open reading frame, but the presence of premature termination codons cause EJCs to remain within aberrant 3’ UTRs, thereby triggering NMD. However, as introns do not inherently elicit NMD in *Drosophila*, its pathway does not appear to involve the EJC (Nicholson and Muhlemann 2010).

The EJC impacts other aspects of the transcript lifecycle, including PolII promoter-proximal pause release (Akhtar et al. 2019), mRNA localization (Le Hir et al. 2001; Palacios et al. 2004) and translation (Wiegand et al. 2003). In particular, given central connections between the EJC and the spliceosome (Singh et al. 2012), attention has been paid to splicing-related functions of the EJC. In *Drosophila*, the EJC positively regulates splicing of long introns, such as *mapk* (Ashton-Beaucage et al. 2010; Roignant and Treisman 2010), and also activates suboptimal splice sites, such as within *piwi* (Hayashi et al. 2014; Malone et al. 2014). By contrast, recent analysis of the mammalian EJC shows that many of its direct splicing targets are instead inhibited. In particular, the Gehring and Ule groups found that deposition of the EJC directly prevents usage of spurious exonic 5’ and 3’ splice sites (Blazquez et al. 2018; Boehm et al. 2018).

With these diverse and often opposing roles in mind, we analyzed the effects of the *Drosophila* EJC on splicing in greater detail. Although *Drosophila melanogaster* has one of the best annotated metazoan transcriptomes (Brown et al. 2014; Westholm et al. 2014; Sanfilippo et al. 2017), we unexpectedly detect many hundreds of novel splice junctions upon depletion of core EJC components in a single celltype. As in mammals, the fly EJC protects neighboring introns from cryptic splice site activation by occlusion, and this function is required at weak splice sites and under unusual circumstances including out-of-order splicing and recursive splicing. Surprisingly, we discover that a subset of novel junctions arise from resplicing within transcript segments that have already undergone intron removal. These may reflect an intrinsic requirement of the EJC to suppress regenerated splice sites that might otherwise undergo recursive splicing. This suggests two distinct stages of EJC protection: (1) suppression of cryptic splice sites during pre-mRNA processing and (2) protection of mRNAs from further resplicing after intron removal. Overall, our findings expand a newly appreciated, ancestral function of the EJC, and emphasize that bypass of this regulatory process via cDNA constructs can have unexpected deleterious consequences.

## Results and Discussion

### EJC depletion leads to activation of spurious junctions

Recently, Roignant and colleagues reported RNA-seq datasets from S2 cells depleted for core EJC factors *eIF4AIII, tsu* (Y14) and *mago* (Akhtar et al. 2019). We re-examined these data for splicing defects, and paid particular attention to spurious splice site usage. We utilized MAJIQ to acquire currently unannotated junctions (3606 novel splice sites supported by ≥5 split reads in the aggregate data), of which 1677 were >2-fold upregulated in at least one EJC-KD condition. As the three core EJC factors are mutually required for stable EJC association at exon-exon junctions, we might expect these to reveal a set of common molecular defects. Indeed, there was both substantial and significant overlap in novel junctions amongst all three conditions (*p-value* < 1×10^−8^ for three-way overlap), and 876 junctions were elevated in two out of three EJC-KD datasets (**Figure 1 - figure supplement 1A**). To introduce further stringency, we also filtered for >2-fold PSI change in 2/3 EJC depletions, yielding 573 spurious junctions from 386 genes (**Figure 1 - figure supplement 1B** and **Supplementary Table 1)**. These genes are diverse, with gene ontology (GO) analysis comprising diverse cellular processes including system development and signaling (**Supplementary Table 2**).

The most frequent spurious junctions involved activation of exonic, alternative 5’ or 3’ SS, followed by novel alternative splicing and intronic SS activation (**Figure 1A**). These are expected to delete exonic sequence (alternative 5’ or 3’ SS) or insert intronic sequence (intronic SS), relative to canonical mRNA products. We depict *straw* as an example of aberrant splicing occuring at a constitutive exon-exon junction (**Figure 1B**). Here, depletion of *eIF4AIII, tsu* and *mago*, but not *lacZ* control, all induced high-frequency usage of a novel exonic, alternative 5’ SS that joins to the constitutive 3’ SS 3248 nt downstream. Importantly, this presumably defective transcript comprises the major isoform in all three core-EJC knockdowns, as it removes 91 nt of coding sequence and is thus out of frame.

**Figure 1.**
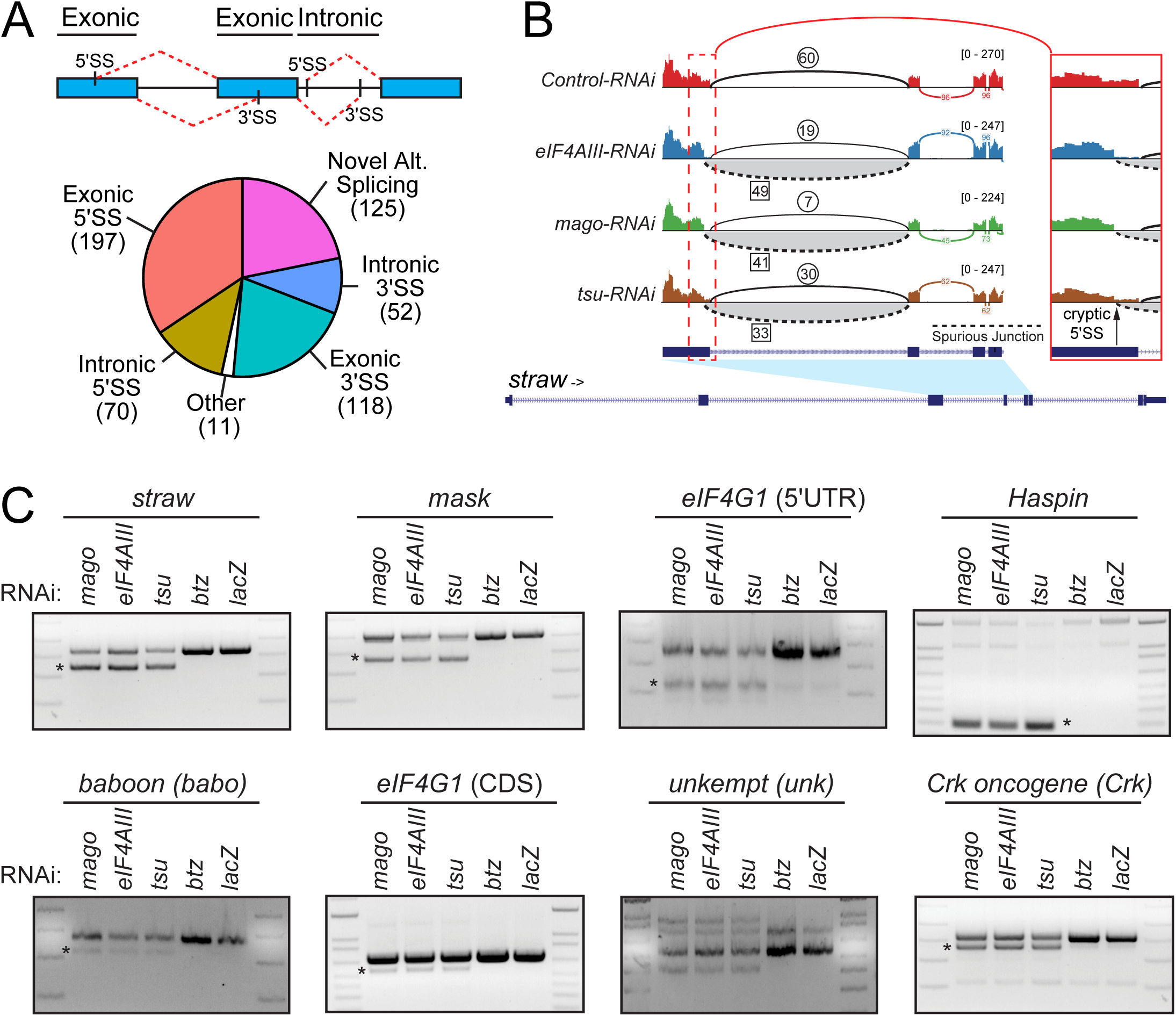
Transcriptome-wide *de novo* alternative splicing upon depletion of functional Exon Junction complex. (A) Overview of upregulated *de novo* splice junctions in EJC-depleted cells. Top: schematic of exonic and intronic cryptic 5’ and 3’ SS. Bottom: Pie chart indicating the distribution of different splice junction classes. (B) Sashimi plot depicting HISAT2-mapped sequencing coverage along a portion of *straw*, which has defective splicing under core-EJC LOF. The gene model depicts the location of the cryptic 5’ SS relative to the annotated 5’ SS. Junction spanning read counts mapping to the canonical junction are circled, whereas cryptic junction read counts are squared. Note that spliced reads mapping to the cryptic junction are found in *eIF4AIII-, mago-* and *tsu-*KD but not the control comparison. Region containing the cryptic 5’SS has been zoomed on the right. (C) Validation of *de novo* splicing events in core-EJC depleted cells. EJC core components (*eIF4AIII, mago, tsu* and *btz*) were knocked down in *Drosophila* S2 cells using dsRNA. After knockdown, eight targets identified in (A) were evaluated using an rt-PCR assay and demonstrated splicing defects (asterisk). Importantly, only core-EJC factor KD produced cryptic bands, but not *btz* or control conditions. Note that several splicing defects are observed for *unkempt (unk)*.

We used rt-PCR to validate *de novo* splice isoforms in EJC-depleted S2 cells. We selected transcripts with high activation of exonic 5’ and 3’ SS (PSI > 0.2), such as *straw, multiple ankyrin repeats single KH domain (mask), baboon* and *eukaryotic translation initiation factor 4G1 (eIF4G1)*, but also evaluated targets with moderate changes (0.01< PSI < 0.05) such as *Crk oncogene and unkempt*. As EJC stabilization during pre-mRNA processing requires *eIF4AIII, tsu* and *mago*, but not *btz*, we utilized knockdown of *btz* and *lacZ* as controls (**Figure 1 - figure supplement 1C**). For all eight amplicons tested, we observed splicing defects only under core-EJC (*eIF4AIII, tsu* and *mago*) knockdown conditions (**Figure 1C**). These data provide stringent validation of our annotation of spurious junctions, and highlight a previously unappreciated quality control function of the *Drosophila* EJC.

### The EJC suppresses cryptic exonic 3’ SS during pre-mRNA processing

These alterations in transcript processing were reminiscent of how the human EJC, recruited to exon junctions, directly influences the splicing of neighboring introns (Blazquez et al. 2018; Boehm et al. 2018). Accordingly, we examined the mechanism of EJC-regulated splicing defects in *Drosophila*. We began by examining transcripts with spurious exonic 3’ SS. These represent a majority of *de novo* events observed in our analysis, and are predicted to cause broad loss of mRNA sequences. Cryptic 3’ SS exhibit strong positional bias and cluster specifically around exon junctions **(Figure 2A**). However, while cryptic 3’ SS contain the invariant 3’ AG dinucleotide (**Figure 2 - figure supplement 1A**), quantitative assessment of SS strength indicated broad variation (**Figure 2A**). In fact, most activated 3’ SS in this category are extremely weak and would not normally be considered functionally competent, especially when considering their sheer frequency in the transcriptome at large. Thus, it was important to manipulate these RNA substrates to understand their splicing capacities more directly.

**Figure 2.**
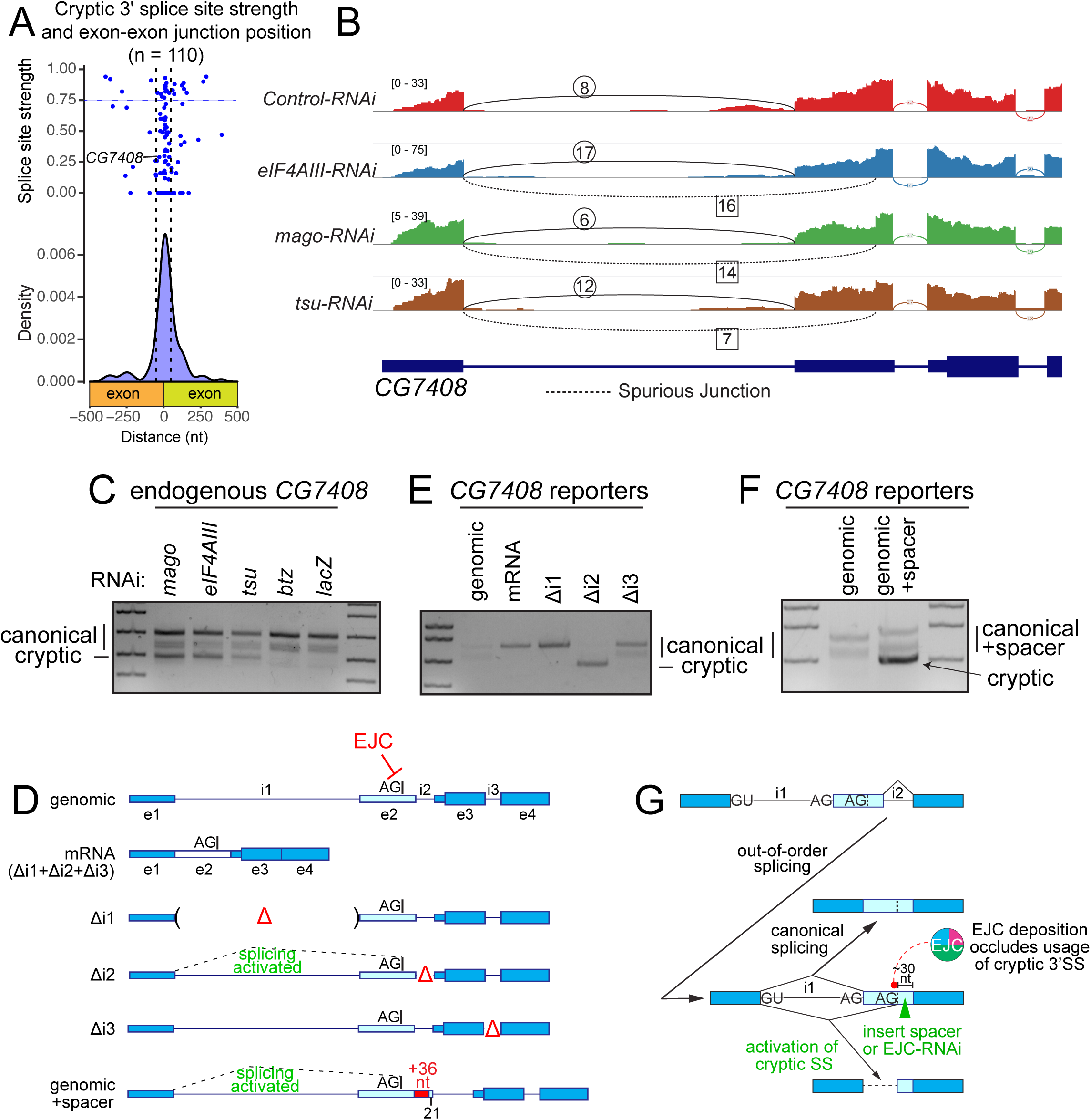
EJC-depletion leads to activation of cryptic 3’ splice sites. (A) Depiction of 3’ SS position of spurious junctions relative to exon-exon boundaries as density and dot plot. The dot plot indicates splice site scores as calculated via NNSPLICE. Horizontal dashed line depicts threshold for strong 3’ SS, and vertical dashed lines specify 50 nt flanking exon-exon junctions. (B) Sashimi plot depicting HISAT2-mapped sequencing coverage along a portion of *CG7408*, which has a cryptic 3’ SS that is activated under core-EJC LOF. Junction spanning read counts mapping to the canonical junction are circled, whereas cryptic junction read counts are squared. Note that spliced reads mapping to the cryptic junction are found in *eIF4AIII, mago* and *tsu* KD but not the control comparison. (C) Validation of *CG7408* cryptic 3’ SS activation in core-EJC, but not *btz* or *lacZ* KD conditions (D) Schematic of *CG7408* splicing reporters. Exons 1-4 (introns included) were cloned and subjected to further manipulation. Locations of pre-removed introns (Δ), as well as a construct lacking all introns (mRNA) are included. For reference, the position of the cryptic 3’SS is marked on exon 2. genomic+spacer represents a modified version of the genomic splicing reporter with an insertion of 36 nt spacer sequence on exon 2. (E) rt-PCR of reporter (D) constructs ectopically expressed in S2 cells demonstrates that intron 2 is required for accurate processing of the minigene. Canonical and cryptic products are indicated. (F) Cryptic splicing is detected with the inclusion of a 36 nt spacer sequence. (G) Schematic of out-of-order splicing and positional requirement of the core-EJC for accurate 3’ SS definition.

We selected *CG7408* as a paradigm: it reproducibly exhibited defective splice isoforms in all core-EJC knockdowns (**Figure 2B**), but its putative 3’ SS is extremely weak (NNSPLICE score of 0.29, **Figure 2A**) and poorly conserved (**Figure 2 - figure supplement 1B**). We used rt-PCR to validate the expected transcript defects in EJC-depleted cells (**Figure 2C)**, and confirmed 183 nt exon deletion relative to the canonical splice isoform via Sanger sequencing. Our cryptic junction replaces intron 1, where canonical splicing typically utilizes one of three annotated 3’ SS, the dominant of which is stereotypically strong (**Figure 2 - figure supplement 1B**, NNSPLICE score of 0.91). We then constructed a minigene bearing exons 1-4 of *CG7408* (**Figure 2D**, genomic). When transfected into S2 cells, this reporter recapitulated normal splicing through activation of annotated 3’ SS (**Figure 2E**, genomic). Importantly, a “fully pre-spliced” reporter lacking all introns, i.e., mimicking an mRNA expression construct, yielded a single normal product (**Figure 2D-E**, mRNA). Thus, pre-processed *CG7408* transcripts that cannot recruit EJC, also do not undergo further processing. At face value, this appears consistent with the hypothesis that the EJC regulates splicing of flanking introns.

We explored this further by testing for potentially distinct consequences of EJC recruitment to individual *CG7408* exon junctions, by removing each intron in turn (**Figure 2D** - Δi1, Δi2 and Δi3). These manipulations should only abolish EJC recruitment at individual pre-processed exon junctions. Δi1 only produced the dominant canonical isoform and Δi3 produced the two known canonical isoforms at the same proportions as the genomic construct (**Figure 2E**). By contrast, pre-removal of intron 2 yielded fully aberrant transcripts (**Figure 2E**, Δi2). These tests emphasize the functional requirement of intron 2 for correct processing of *CG7408* and demonstrate that even poor matches to consensus splice sites (i.e., the *CG7408* cryptic 3’ SS) can be potently activated in the absence of the EJC.

We emphasize that these data support a mechanism in which intron 2 is excised first, and this order is required for the correct definition of the annotated intron 1 3’ SS (**Figure 2G**). Out-of-order splicing has been previously observed (Takahara et al. 2002; Drexler et al. 2020), but its relationship to accurate pre-mRNA maturation has generally been unclear. These experiments, along with recent work by Gehring and colleagues (Boehm et al. 2018) indicate a requirement for out-of-order splicing for proper mRNA maturation.

How does the EJC inhibit definition of cryptic exonic 3’ SS? In human cells, the EJC can directly mask cryptic 3’ SS. Based on the close clustering of these sites around the position of EJC recruitment (**Figure 2A**), we reasoned that the EJC may occlude important features of the 3’ SS, such as the branchpoint, polypyrimidine tract or 3’ intron junction that base-pairs with the U2 snRNP complex. We tested this hypothesis by separating the cryptic 3’ SS on our genomic reporter from the site of EJC recruitment, by inserting a 36 nt spacer (**Figure 2D**, genomic+Spacer). Unlike the genomic construct, which yields only annotated splice isoforms, the genomic+Spacer variant yielded additional truncated transcripts, consistent with derepression of the cryptic 3’ SS (**Figure 2F**). Altogether, these data demonstrate the fly EJC aids accurate SS selection during pre-mRNA processing by masking cryptic 3’ SS.

### The EJC prevents cryptic exonic 5’ SS activation during pre-mRNA processing

We next used analogous strategies to study cryptic exonic 5’ SS. These sites represent ∼35% of novel splice junctions upregulated under EJC-depleted conditions and are expected to be deleterious to mRNA processing fidelity. Bioinformatic analysis indicated that cryptic 5’ SS share general structural properties with 3’ SS, such as clear preference in the vicinity of exon junctions but distribution across a wide range of strengths (**Figure 3A, Figure 3 - figure supplement 1A**).

**Figure 3.**
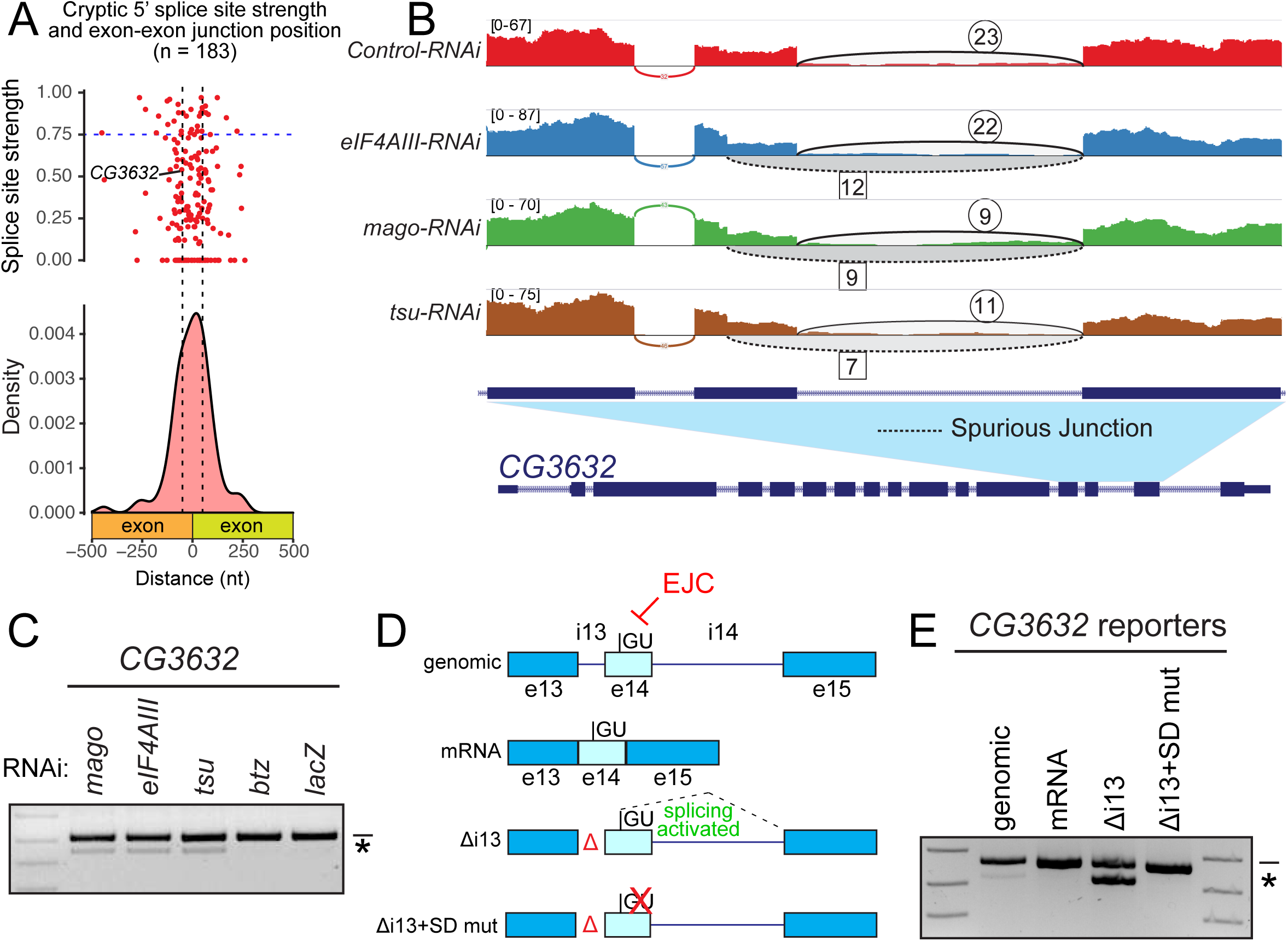
EJC-depletion leads to activation of cryptic 5’ splice sites. (A) Metagene of cryptic 5’ SS position relative to exon-exon boundaries as density and dot plot. The dot plot indicates splice site scores as calculated via NNSPLICE (see Methods). Horizontal dashed line depicts threshold for strong 3’ SS, and vertical dashed lines specify 50 nt flanking exon-exon junctions. (B) Sashimi plot depicting HISAT2-mapped sequencing coverage along a portion of *CG3632*, which has a cryptic 5’ SS that is activated under core-EJC LOF. Junction spanning read counts mapping to the canonical junction are circled, whereas cryptic junction read counts are squared. Note that spliced reads mapping to the cryptic junction are found in *eIF4AIII, mago* and *tsu* KD but not the control comparison. (C) Validation of *CG3632* cryptic 5’ SS activation (asterisk) in core-EJC, but not *btz* or *lacZ* KD conditions (D) Schematic of *CG3632* splicing reporters. Exons 13-15 (introns included) were cloned and subjected to further manipulation. Locations of pre-removed introns (Δ), as well as a construct lacking all introns (mRNA) are included. The position of the cryptic 5’SS is marked on exon 14, and was mutated in Δi13+SD mut. (E) rt-PCR of reporter (D) constructs ectopically expressed in S2 cells demonstrates that intron 13 is required for accurate processing of the minigene. Canonical products are indicated by the line and cryptic products by an asterisk.

We selected *CG3632* for mechanistic tests, as core-EJC knockdown data showed activation of a poorly conserved, weak cryptic 5’ SS (**Figure 3B** and **Figure 3 - figure supplement 1B-C** – NNSPLICE score of 0.54) on exon 14. Using rt-PCR and Sanger sequencing, we validated that EJC-depletion induces a defective *CG3632* splice isoform lacking 71 nt of coding sequence (**Figure 3C**).

We hypothesized that the EJC, recruited to the exon 13/14 junction, suppresses the cryptic 5’ SS on exon 14 and activates the canonical 5’ SS during removal of intron 14. We tested this using a minigene reporter consisting of exon 14 (containing the cryptic 5’ SS) and its immediately flanking introns and exons (**Figure 3D**, genomic). Expression of this reporter in S2 cells predominantly resulted in the canonical product, but we also observed a minor amount of cryptic 5’ SS activation (**Figure 3E**, genomic). As a negative control, we generated a version lacking both introns (**Figure 3D**, Δi13+14), which produced the expected mRNA (**Figure 3E**, Δi13+14). Notably, removal of intron 13 alone (**Figure 3D**, Δi13), mimicking loss of EJC recruitment at the exon 13/14 junction, yielded high levels of cryptic 5’ SS activation (**Figure 3E**, Δi13) that were fully suppressed by mutation of the cryptic 5’ SS in the Δi13 reporter (**Figure 3D-E**, Δi13+SD mut). Altogether, these data support that deposition of the EJC during pre-mRNA processing suppresses cryptic 5’ SS during subsequent intron removal.

### The EJC suppresses recursive splice sites

Given that the EJC suppresses both 5’ and 3’ SS, a potentially more complex scenario might exist if both types of cryptic splice sites were to be activated in the vicinity of each other. We inspected our catalog of spurious junctions for this possibility, and considered that even modest matches to consensus splice sites (**Figure 2A and 3A**) might serve as viable candidates for further evaluation. Interestingly, many exon junctions were potentially able to regenerate weak splice sites after intron removal, reminiscent of the process of recursive splicing (RS) (Burnette et al. 2005).

We first investigated a spurious junction within *Casein kinase IIß* (*CkIIß*), where core-EJC LOF led to loss of 54 nt of canonical mRNA sequence (**Figure 4 - figure supplement 1A-B**). Assessment of the novel 3’ SS on exon 3 revealed that it lacks a polypyrimidine tract and is a poor match to the consensus (**Figure 4A**). On the surface, the mechanism of cryptic 3’ SS activation on *CkIIβ* might appear similar to that of *CG7408* (**Figure 2F, Figure 4 - figure supplement 1D, path 2**). However, upon examining *CkIIβ* for splice sites, we found an additional poor recursive 5’ SS at the beginning of exon 3 (**Figure 4A**). Therefore, we imagined an alternate scenario, whereby dual cryptic 5’ and 3’ SS might be derepressed upon EJC loss, leading to resplicing (**Figure 4 - figure supplement 1D, path 1**). Crucially, whether one-step splicing (via alternative splicing) or resplicing (via recursive splicing event), the resulting mRNA products are indistinguishable (**Figure 4A**). Therefore, we devised reporter tests to clarify the underlying mechanism.

**Figure 4.**
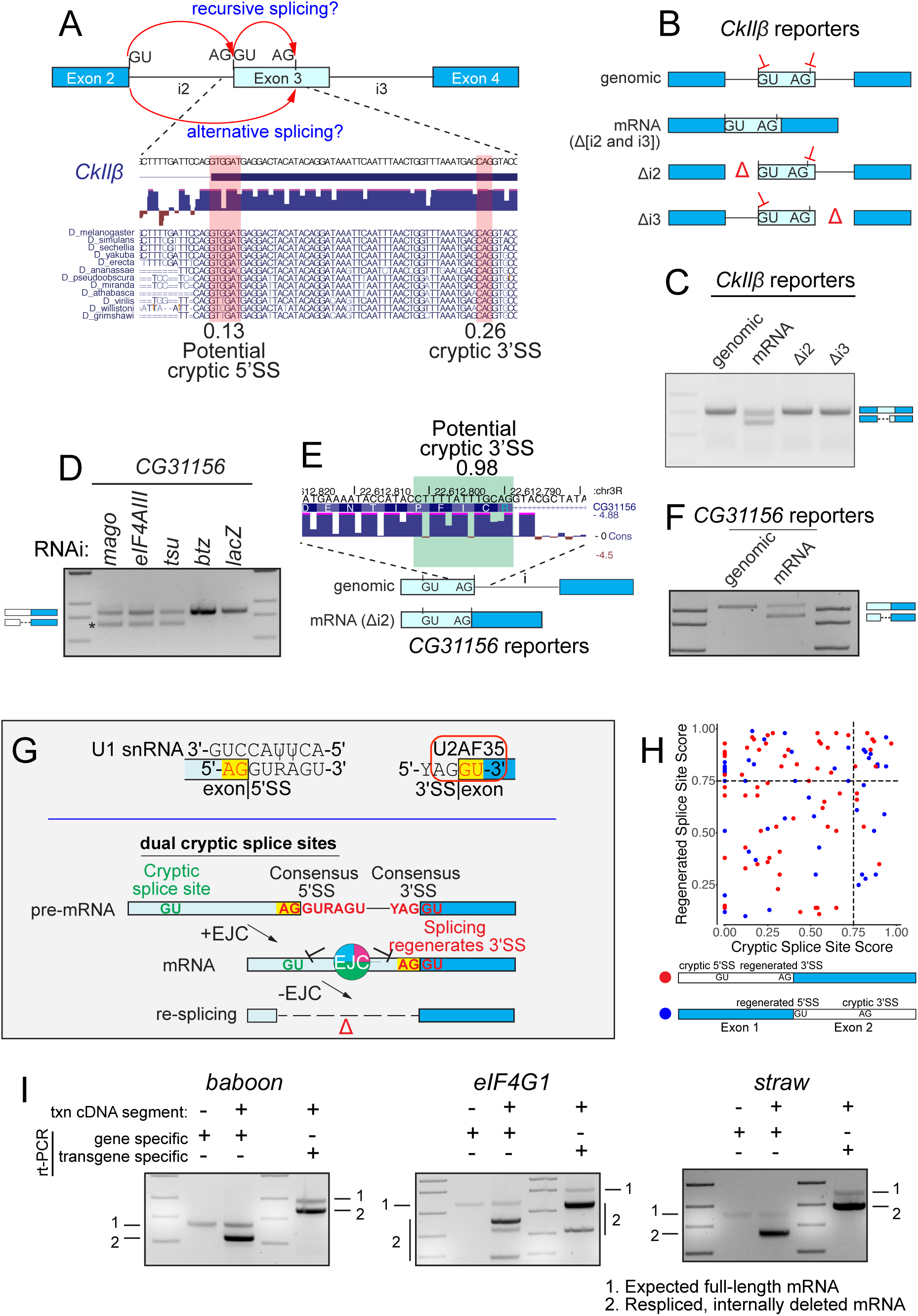
EJC-depletion leads to activation of dual cryptic splice sites and resplicing of mRNA. (A) Above: Schematic of resplicing splicing versus alternative resplicing, both of which would yield the same aberrant mRNA product. Below: Sequence of *CkIIβ* transcript lost due to cryptic splicing. Cryptic 3’ SS activated is highlighted in red, as well as a potential regenerated 5’ SS. Scores listed are generated by NNSPLICE. Conservation across Drosophilid family is shown. (B) Schematic of *CkIIβ* splicing reporters. Exons 2-4 (introns included) were cloned and subjected to further manipulation. Locations of pre-removed introns (Δ), as well as a construct lacking all introns (mRNA) are included. For reference, the position of the cryptic 3’ SS and potential 5’ recursive splice sites is marked on exon 3. (C) rt-PCR of *CkIIβ* reporter constructs in S2 cells demonstrates that introns are required for accurate processing of the minigene. Canonical and cryptic products are indicated. (D) Validation of *CG31156* cryptic 5’ SS activation in core-EJC, but not *btz* or *lacZ* KD conditions (E) Schematic of *CG31156* splicing reporters with and without introns. Location of potential 3’ recursive splice site on exon 2 is indicated along with conservation scores. (F) rt-PCR of reporter constructs in S2 cells demonstrates that introns are required for accurate processing of the minigene. Canonical and cryptic products are indicated. (G) Model for mRNA resplicing. Top, Binding sites of U1 snRNA and U2AF35 define the 5’ SS and 3’ SS, respectively, but also impose constraints on flanking exonic sequences that intrinsically regenerate splice site mimics in a recursive fashion. (Bottom) When located in proximity to another cryptic splice site, these can lead to mRNA resplicing in the absence of the EJC. An example of dual cryptic splice sites with a regenerated 3’ SS is shown, but this can also occur with a regenerated 5’ SS. (H) Comparison of splice site strengths for cases of dual cryptic splice site activation. Cases that contain regenerated 3’ and 5’ splice sites at exon junctions and their structures are schematized and distinguished by red and blue dot. Dashed lines mark thresholds for reasonably strong splice sites. (I) Resplicing on cDNAs. Constructs bearing cDNA segments of *baboon, eIF4G1* and *straw* were expressed in S2 cells and yielded re-spliced amplicons. Gene specific primers that amplify endogenous and ectopic products only show resplicing from the intron-less reporter. Transgene-specific primers demonstrate mostly re-spliced products.

We first used rt-PCR to validate that core-EJC knockdown resulted in substantial activation of a truncated *CkIIß* splice isoform corresponding to RNA-seq data (**Figure 4 - figure supplement 1C**). We then analyzed a series of splicing minigenes (**Figure 4B**). Expression of *CkIIβ* exons 2-4 with all introns present produced a single product with the expected introns spliced out (**Figure 4C**, genomic). We precisely tested the positional necessity of the EJC at each exon junction by pre-removing each intron (**Figure 4B**, Δi2 and Δi3). These reporters also underwent normal splicing (**Figure 4C**, Δi2 and Δi3), demonstrating that *CkIIβ* processing defects were in fact mechanistically distinct from those determined for *CG7408*. Strikingly, upon testing a construct with both introns pre-removed, we observed a switch to truncated product output, corresponding to activation of the unannotated recursive 5’ SS and 3’ SS (**Figure 4B-C**, mRNA). This supports a model where the EJC is required at multiple positions to repress spurious 5’ and 3’ SS simultaneously (**Figure 4 - figure supplement 1D, path 1**).

We characterized another instance of dual cryptic splice site within *CG31156*, albeit of a different flavor. Here, sashimi plots indicate activation of an exonic 5’ SS within exon 2 (**Figure 4 - figure supplement 2A-B**) and we validated this 110 nt deletion isoform using rt-PCR (**Figure 4D**). Importantly, based on these data alone, it would be reasonable to predict this as a case of alternative cryptic 5’ SS activation. However, we noticed that removal of the canonical intron 2 regenerates a putative recursive 3’ SS at the exon 2/3 boundary (**Figure 4E, Figure 4 - figure supplement 2C**). Therefore, we examined reporters to examine the mechanism underlying this unwanted splicing pattern. Expression of the genomic reporter that required intron removal yielded the expected mRNA product (**Figure 4F**, genomic). Conversely, pre-removal of the intron and expression of the mRNA resulted in the truncated re-spliced product (**Figure 4F**, mRNA). Accordingly, these data again indicate that the EJC represses dual cryptic splice sites during mRNA processing (**Figure 4 - figure supplement 2D**).

Previous annotations of *Drosophila* recursive splice sites (RSS) concluded that these hybrid 5’/3’ SS are highly conserved, flanked by short cryptic downstream exons, and are highly biased to reside in long introns (mean length ∼50 kb) (Duff et al. 2015; Joseph et al. 2018). While it is possible that recursive splicing aids processing of long introns, it is also conceivable that it is easier to capture RS intermediates within long introns. The examples of cryptic RSS on the *CkIIβ* and *CG31156* transcripts clearly deviate from canonical RSS architectural properties, i.e., they are hosted in short introns and exhibit modest to poor conservation. Moreover, the example of a recursive 3’ SS in *CG31156* is to our knowledge the first validated instance, and represents a conceptually novel RSS location. Importantly, the relevant AG dinucleotide in the *CG31156* 3’ recursive splice site is not preserved beyond the closest species in the melanogaster subgroup (**Figure 4 - figure supplement 2C**), and the amino acids encoded by the functional 5’ RSS in *CkIIβ* diverge with clear wobble patterns (**Figure 4A**). Thus, these examples of cryptic exonic recursive splicing are functional, but evolutionarily fortuitous.

### The EJC protects spliced mRNAs from resplicing

Since many genes span large genomic regions, cDNA constructs have been a mainstay of directed expression strategies. It is generally expected that these should be effective at inducing gain-of-function conditions, yet cDNA constructs are not typically vetted for proper processing. Our finding of dual cryptic splice sites on transcripts was alarming because in both cases, we observed resplicing on mRNA constructs (**Figure 4C and 4F**). To reiterate, the EJC prevents dual cryptic SS from resplicing on transcript segments that have already undergone intron removal, but such protection will be missing from intronless cDNA copies.

We were keen to assess the breadth of this concept. To do so, we examined the sequence of mRNAs bearing EJC-suppressed cryptic SS, and looked for additional unidentified, complementary SS. Notably, since resplicing would have to map to a canonical junction, we looked for regenerated SS at exon junctions. An initial survey for SS invariant dinucleotide signatures (AG for 3’ SS and GT for 5’ SS) indicated that 64/118 junctions with cryptic 3’ SS and 104/183 junctions with cryptic 5’ SS were compatible with resplicing. The fact that over half of both classes of cryptic splicing events were potentially compatible with resplicing might at first glance seem like a tremendous enrichment. However, it does in fact reflect fundamental features of extended consensus splice sequences that basepair with the spliceosome, namely the U1 snRNP and U2AF35 binding sites, respectively (**Figure 4G**-top). Quantification of these sequences indicated a range of regenerated 5’ and 3’ SS at exon junctions, with at least 59 junctions resembling strong SS (**Figure 4H**, NNSPLICE>0.75). However, as several cryptic 5’ and 3’ SS amongst our validated loci (**Figures 1-4**) were extremely poor, with functional dual cryptic splice sites in *CkIIβ* scoring at only 0.13 and 0.26 (**Figure 4A**), the functional breadth of this phenomenon is undoubtedly broader. Therefore, we imagined a scenario where a core function of the EJC is to repress splice sites that were regenerated at exon junctions as a consequence of intron removal using canonical splice sites (**Figure 4G**-bottom).

Nevertheless, as this model cannot be explicitly distinguished from alternative splicing without experimental testing, we selected additional loci for analysis. Therefore, we constructed partial cDNA constructs for three genes, encompassing regions we had validated as subject to EJC-suppression of cryptic splicing (**Figure 1C**), and selected targets that survey a range of regenerated SS strengths. These include *straw*, which yields a strong 3’ RSS (NNSPLICE score of 0.98) after removal of intron 3; *eIF4G1*, which regenerates a moderate 5’ RSS (NNSPLICE score of 0.64) after processing of intron 10; and *baboon*, which produces an exceptionally poor 3’ RSS (NNSPLICE score of 0) after removal of intron 4, bearing only the AG dinucleotide.

In contrast to the endogenous genes which produced a single amplicon, expression of all three cDNA constructs yielded substantial re-spliced products, supporting our view that the EJC prevents activation of dual cryptic SS on mRNAs, including SS regenerated at exon junctions (**Figure 4I**). Unexpectedly, SS strength did not correlate with levels of resplicing. Indeed, the majority of transcripts from all three reporters were truncated, including from *baboon*. Furthermore, the *eIF4G1* reporter yielded three truncated products, suggesting that other sequences may also serve as cryptic SS. As these examples of resplicing occur on coding regions of the transcript, all of them either delete amino acids or generate frameshifts. We conclude that many cDNA constructs are potentially prone to resplicing due to loss of protection afforded by the EJC.

### Conserved role for the EJC to repress cryptic splicing and implications for cDNA expression

Although introns are not essential for gene expression, they play important facilitatory roles by enhancing export and translation in part through recruitment of the EJC during splicing. Subsequently, it was recognized that once deposited, the EJC also promotes accurate gene expression by regulating processing of neighboring introns. Recently, in the mammalian setting, the role of the EJC during pre-mRNA splicing was extended to include suppression of cryptic splice sites (Blazquez et al. 2018; Boehm et al. 2018).

Here we reveal that the fly EJC similarly plays a broad role in direct suppression of cryptic exonic splice sites, owing to its characteristic deposition 20-24 nt upstream of exon-exon junctions. Thus, we now appreciate that concealment and suppression of cryptic splice sites is a conserved EJC activity (Boehm et al. 2018). Importantly, the positional recruitment of the EJC during splicing is conserved and sequence-independent (Le Hir et al. 2000). Thus, we infer this function should also be independent of splice site divergence between phyla, as well as splice site strength, and should not require accessory components. In contrast, non-conserved roles of the EJC appear to rely on integration within and diversification of distinct functional networks. For example, while the Upf (*Up-f*rameshift) proteins coordinate NMD across eukaryotes (He and Jacobson 2015), the mechanisms differ. In mammals, NMD is coordinated with intron removal through direct interactions between the EJC and Upf3 (Kim et al. 2001; Le Hir et al. 2001; Lykke-Andersen et al. 2001). However, these interactions are not found in invertebrates, and consequently the invertebrate EJC is not involved in NMD (Nicholson and Muhlemann 2010).

In addition to pre-mRNAs, we show that the EJC also suppresses cryptic splice sites within spliced mRNAs. Although this mechanism cannot be distinguished from alternative splicing (**Figure 4A**) without further experimentaion, we readily detect resplicing on all cDNA constructs tested. Unexpectedly, while these junctions appear to contain just one cryptic SS, our data indicates that these transcripts contain secondary cryptic splice sites that mediate resplicing. Importantly, we validate that even poor matches to SS consensus motifs are competent for resplicing. Curiously, as all of our demonstrated examples involve a recursive event at either the 5’ or 3’ cryptic SS, our findings broaden a phenomenon that was previously described within long introns (Duff et al. 2015; Joseph et al. 2018). Furthermore, canonical SS sequences that undergo base pairing interactions with U1 snRNA (5’ SS) and U2AF35 (3’ SS) have motifs AG|<u>GURAGU</u> and <u>YAG</u>|GU (Kielkopf et al. 2001; Kondo et al. 2015). It is noteworthy that core splice site signals contain bases that are compatible with regeneration of splice sites and that these naturally occur proximal to EJC recruitment sites. Accordingly, we propose that an ancestral function for the conserved position of EJC deposition may be to prevent accidental activation of regenerated splice sites.

Finally, our observations of resplicing on cDNAs reflect an essential function for introns in protecting mRNA fidelity. For all tested cases of cDNA resplicing on coding sequences, we note deletions of peptide segments or truncations with loss of domains required for protein function (**Figure 4 - figure supplement 3A-C**). Importantly, these affected targets include essential genes, such as *eIF4G1* and activin receptor *baboon*. In the case of *baboon*, the 54 nt splicing defect leads to a deletion of 18 amino acids (195-212, **Figure 4 - figure supplement 3A**). For *eIF4G1*, resplicing removes 131 nt of mRNA sequence, alters the open reading frame and leads to protein truncation with loss of the MI and W2 domains (**Figure 4 - figure supplement 3B**). Finally, resplicing on *straw* transcripts also alters reading frame by removing 91 nt of mRNA, and is predicted to remove 2/3 Plastocyanin-like domains (**Figure 4 - figure supplement 3C**). Thus, our findings have serious implications for functional genomics as well as community genetic studies (Yu et al. 2011; Wei et al. 2020), where cDNA expression constructs and collections are often employed with little attention paid to mRNA processing. Altogether, our work uncovers an important co-transcriptional function of intron removal and the role of the EJC to protect the transcriptome from unwanted resplicing.

## Materials and Methods

### Bioinformatic analysis

The core-EJC knockdown RNA-sequencing datasets were previously reported (Akhtar et al. 2019) and obtained from the NCBI Gene Expression Omnibus (GSE92389). Raw sequencing data was mapped to the *Drosophila* reference genome sequence (BDGP Release 6/dm6) using HISAT2 (Kim et al. 2015) under the default settings. Splice junctions were mapped using the MAJIQ algorithm (2.0) under default conditions (Vaquero-Garcia et al. 2016). Splice graphs and known/novel local splice variants were defined with the MAJIQ Builder using annotations of known genes and splice junctions from Ensembl release 95 and all BAM files. The MAJIQ Quantifier was used to calculate relative abundances (percent selected index - PSI) for all defined junctions. The resulting data was output into tabular format using the Voila function.

A custom R script was written to process all MAJIQ-defined novel junctions relative to the Ensembl gene annotations and identify *de novo* EJC-suppressed junctions. First, we quantified usage of all novel junctions by mining mapped libraries (BAM files) for high quality junction spanning reads with at least 8 nt of overhang and no mismatches. These counts were normalized to sequencing depth per library. To identify *de novo* junctions that may be upregulated, we first selected junctions with at least 5 split reads. In order to enrich for *de novo* junctions that are suppressed by the EJC pathway, we looked for those with > 2 fold difference in at least 2/3 core-EJC RNAi conditions relative to the *lacZ* control. To apply further stringency, we also required that the PSI measurements reflect sufficient change between treatment and control conditions. Therefore, we applied an additional filter of PSI fold change > 2 in at least 2/3 core-EJC RNAi conditions. These criteria produced a total of 573 novel junctions. All junctions are reported in **Supplementary Table 1**.

The 5’ and 3’ ends of these junctions were compared against known gene annotations to characterize splice sites. Exonic 5’ and 3’ SS reflect sites that mapped on exons while the other end mapped to a canonical splice junction, and the same process was used to define intronic 5’ and 3’ SS. *de novo* cases of alternative splicing reflect junctions that utilized annotated splice sites but represented novel connectivity. Sashimi plots were generated using features available on the Integrative Genomics Viewer (IGV) (Robinson et al. 2011).

We generated a custom pipeline to assess recursive splicing potential (Figure 4). Briefly, we identified transcripts that contained cryptic exonic 5’ and 3’ splice sites. For these transcripts, we mapped the position of all splice junctions on the mRNA, which could in theory generate the observed splicing defects. We examined sequences directly downstream of relevant splice junctions to identify potential 5’ recursive splicing and those directly upstream to identify potential 3’ recursive splicing.

We calculated splice site strengths using NNSPLICE (https://www.fruitfly.org/seq_tools/splice.html) (Reese et al. 1997). The sequences used for these analyses were obtained from mRNA rather than the genomic context, which may contain intronic sequences as well. To generate nucleotide content plots, splice sites and their indicated flanking sequences were obtained from mRNAs and fed to WebLogo version 2.8.2 (Crooks et al. 2004). The splice sites are centered in these plots.

All custom scripts used in this study are reported on the Lai lab GitHub page.

### Constructs and cell culture

All splicing reporters were cloned into pAC-5.1-V5-His (ThermoFisher Scientific) using compatible restriction sites. We used PCR to amplify minigene splicing reporters from Drosophila genomic DNA, and used site directed mutagenesis to remove specified introns. We used cDNAs to amplify reporters lacking introns. For genes with multiple isoforms (such as *CG7408*), we cloned the dominant fragment. All primers used for generating constructs and mutagenesis have been summarized in **Supplementary Table 3**.

Transfections were performed using S2-R+ cells cultured in Schneider Drosophila medium with 10% FBS. Cells were seeded in 6-well plates at a density of 1×10^6^ cells/mL and transfected with 200 ng of plasmid using the Effectene transfection kit (Qiagen). Cells were harvested following 3 days of incubation.

### Knockdown of EJC factors in S2 cells

The indicated EJC components were knocked down via RNAi (dsRNA-mediated interference) in S2-R+ cells. The MEGAscript™ RNAi kit (ThermoFisher Scientific) was used to produce dsRNAs required for this experiment. Briefly, DNA templates containing promoter sequences on either 5’ end were produced through PCR with T7-promoter-fused primers. 2 µg of DNA template was transcribed *in vitro* for 4 hours as recommended by the manufacturer. The products were incubated at 75° C for 5 minutes and brought to room temperature to enhance dsRNA formation. A cocktail of DNAseI and RNAse removed DNA and ssRNAs, and the remaining dsRNA was purified using the provided reagents. All dsRNA reagents were verified by running on a 1% agarose gel and quantified by measuring absorbance at 260 nm using a NanoDrop™ (ThermoFisher Scientific).

For knockdown, 3×10^6^ S2-R+ cells in 1 mL serum free medium were incubated with 15 µg of dsRNA for 1 hour at room temperature. Then, 1 mL of medium containing 20% FBS was added to the cells and the whole mixture was moved to a 6 well plate. Cells were collected after 4 days of incubation.

### RT-PCR

After transfection or RNAi treatment, cells were washed in ice cold PBS and pelleted using centrifugation. RNA was collected using the TRIzol reagent (Invitrogen) under the recommended conditions. 5 µg of RNA was treated with Turbo DNase (Ambion) for 45 min before cDNA synthesis using SuperScript III (Life Technology) with random hexamers. RT-PCR was performed using AccuPrime Pfx DNA polymerase (ThermoFisher Scientific) with standard protocol using 26 cycles and primers that were specific to each minigene construct. All primers are listed and described in **Supplementary Table 3**.

## Acknowledgments

Work in E.C.L.’s group was supported by the National Institutes of Health (R01-NS083833 and R01-GM083300) and MSK Core Grant P30-CA008748.

## Author contributions

BJJ designed the experiments, interpreted the data, and helped write the manuscript. ECL helped interpret data and write the manuscript.

## Figure Legends

**Figure 1 - figure supplement 1.**
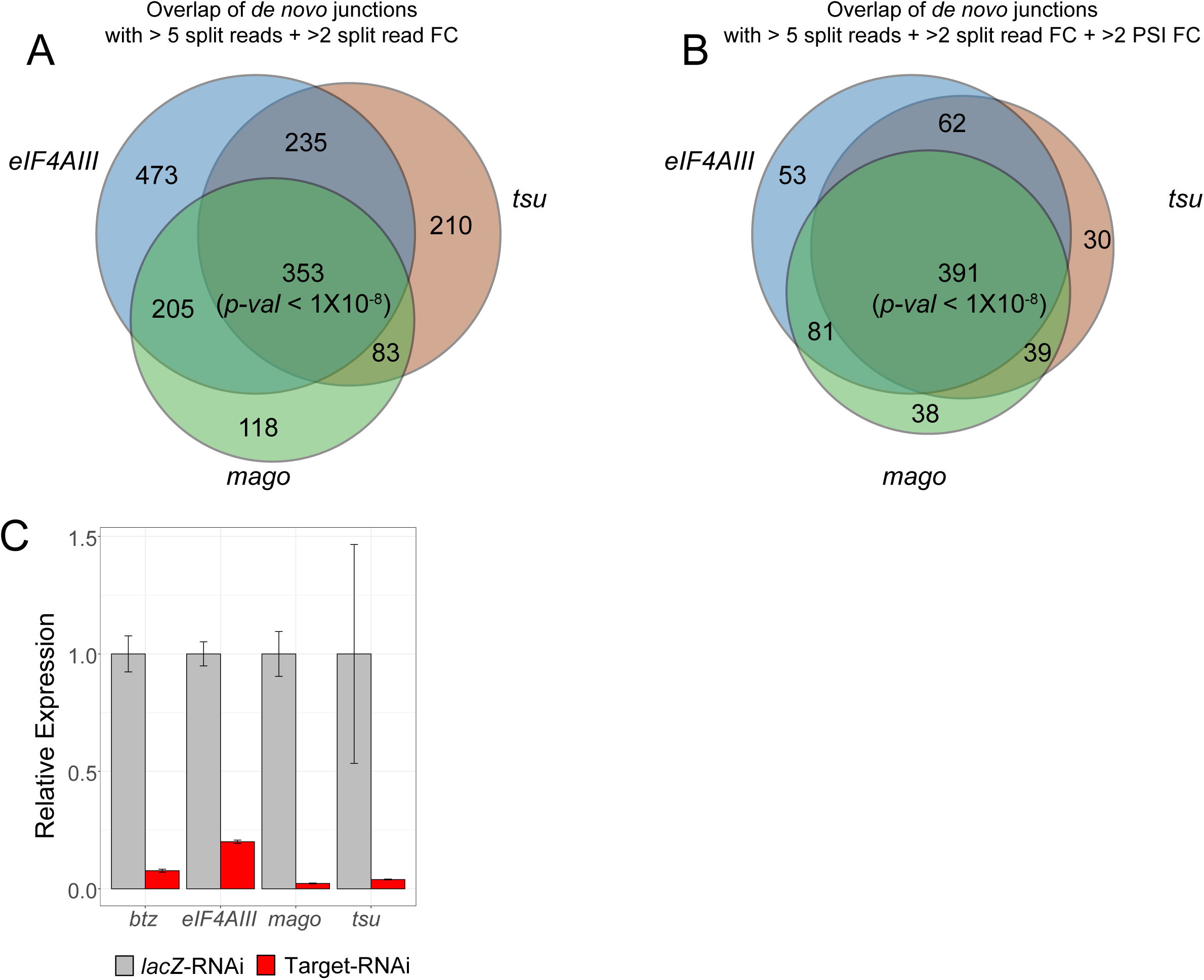
core-EJC depletion yields broad activation of *de novo* splice junctions. (A) Strong overlap of de novo splice junctions between core-EJC knockdown conditions based. The Venn diagram depicts which of 1677 junctions with at least 5 split reads had > 2-fold split read changes between treatment and controls.p-value for three-way overlap was calculated using a permutation test with 10^8^ tests. (B) Strong overlap of high-confidence de novo splice junctions between core-EJC knockdown conditions. The Venn diagram depicts which of 876 junctions with at least 5 split reads and > 2-fold split read changes also show > 2-fold changes in percent selected index (PSI) between treatment and controls. p-value for three-way overlap was calculated using a permutation test with 10^8^ tests. (C) Knockdown of EJC factors in S2 cells using dsRNA. quantitative rt-PCR of core-EJC and btz transcripts after dsRNA treatment.

**Figure 2 - figure supplement 1.**
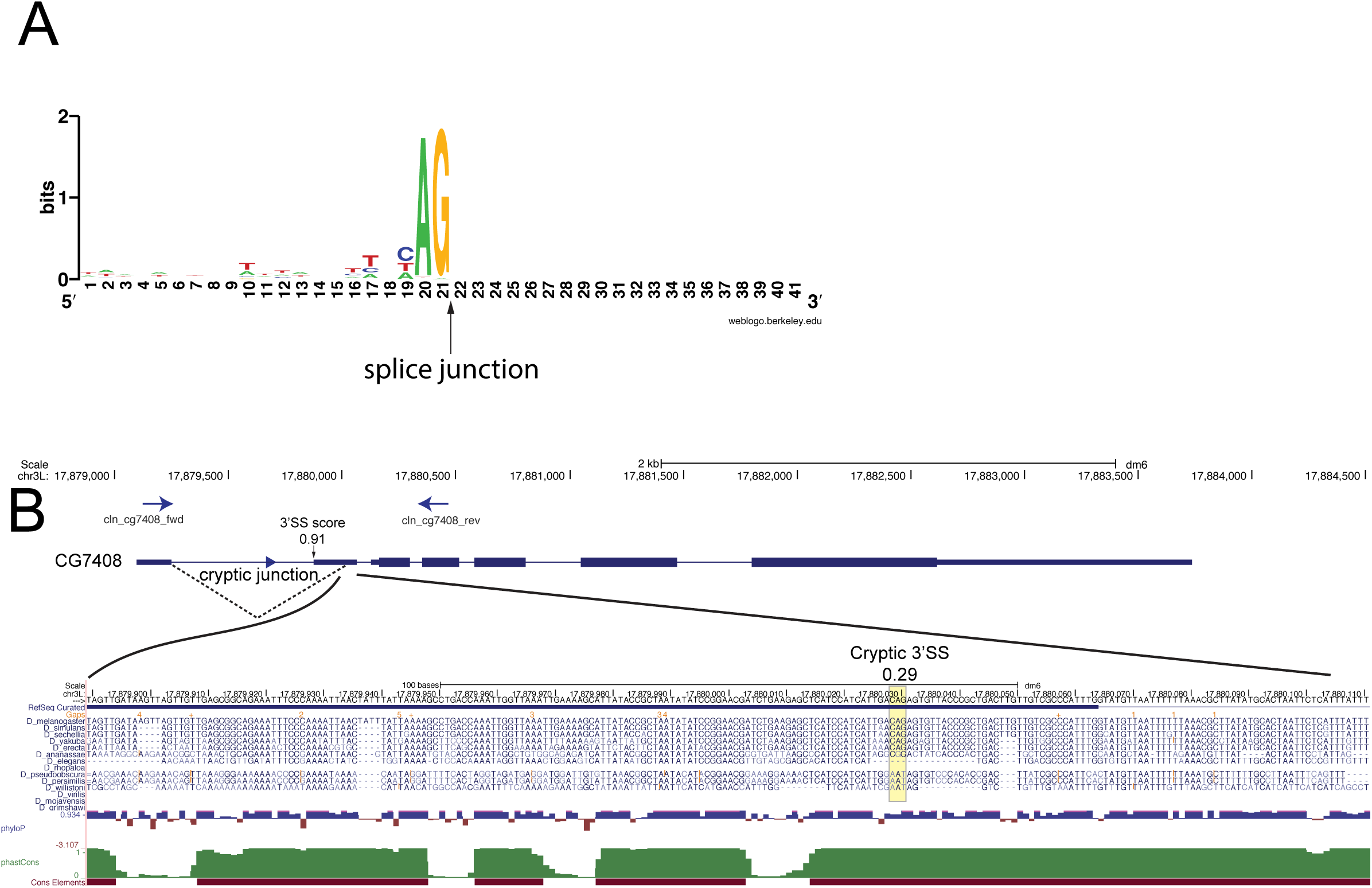
A majority of cryptic 3’ SS activated under EJC-loss are weak. (A) Nucleotide content of cryptic 3’ SS. These sequences, apart from the invariant AG dinucleotide show poor strength. (B) Example of a weak cryptic 3’ SS (NNSPLICE score of 0.29) found on the *CG7408* transcript. (C) Conservation of the weak splice site is depicted using the multiple alignment format on the UCSC genome browser, as well as phyloP and phastCons scores.

**Figure 3 - figure supplement 1.**
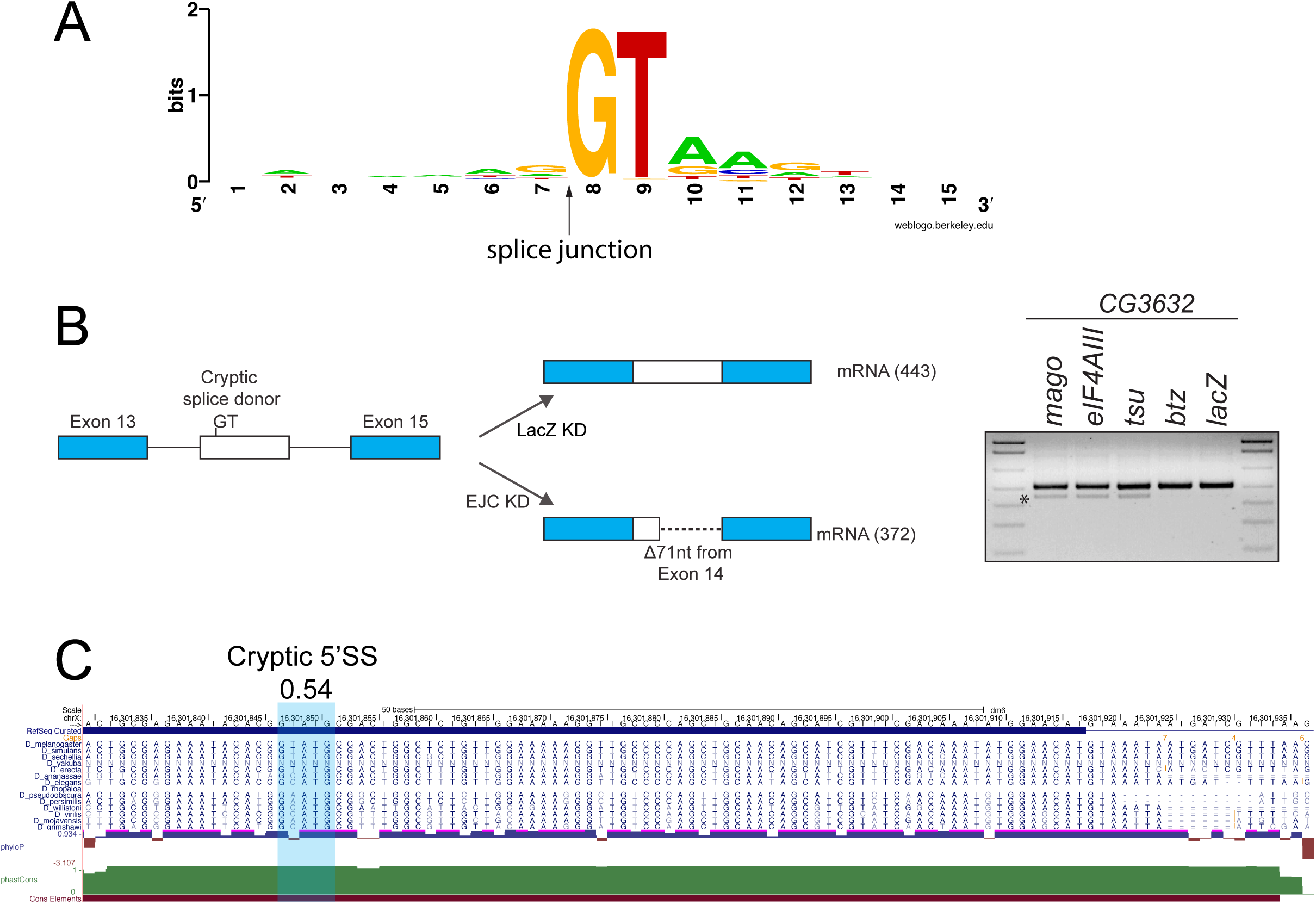
A majority of cryptic 5’ SS activated under EJC-loss are weak. (A) Nucleotide content of cryptic 5’ SS. (B) Schematic of a *de novo* splicing event detected on the *CG3632* transcript. Validation of splicing defects shown on the right. (C) Cryptic 5’ SS (NNSPLICE score of 0.54) found on the *CG3632* transcript. Conservation of the weak splice site is depicted using the multiple alignment format on the UCSC genome browser, as well as phyloP and phastCons scores.

**Figure 4 - figure supplement 1.**
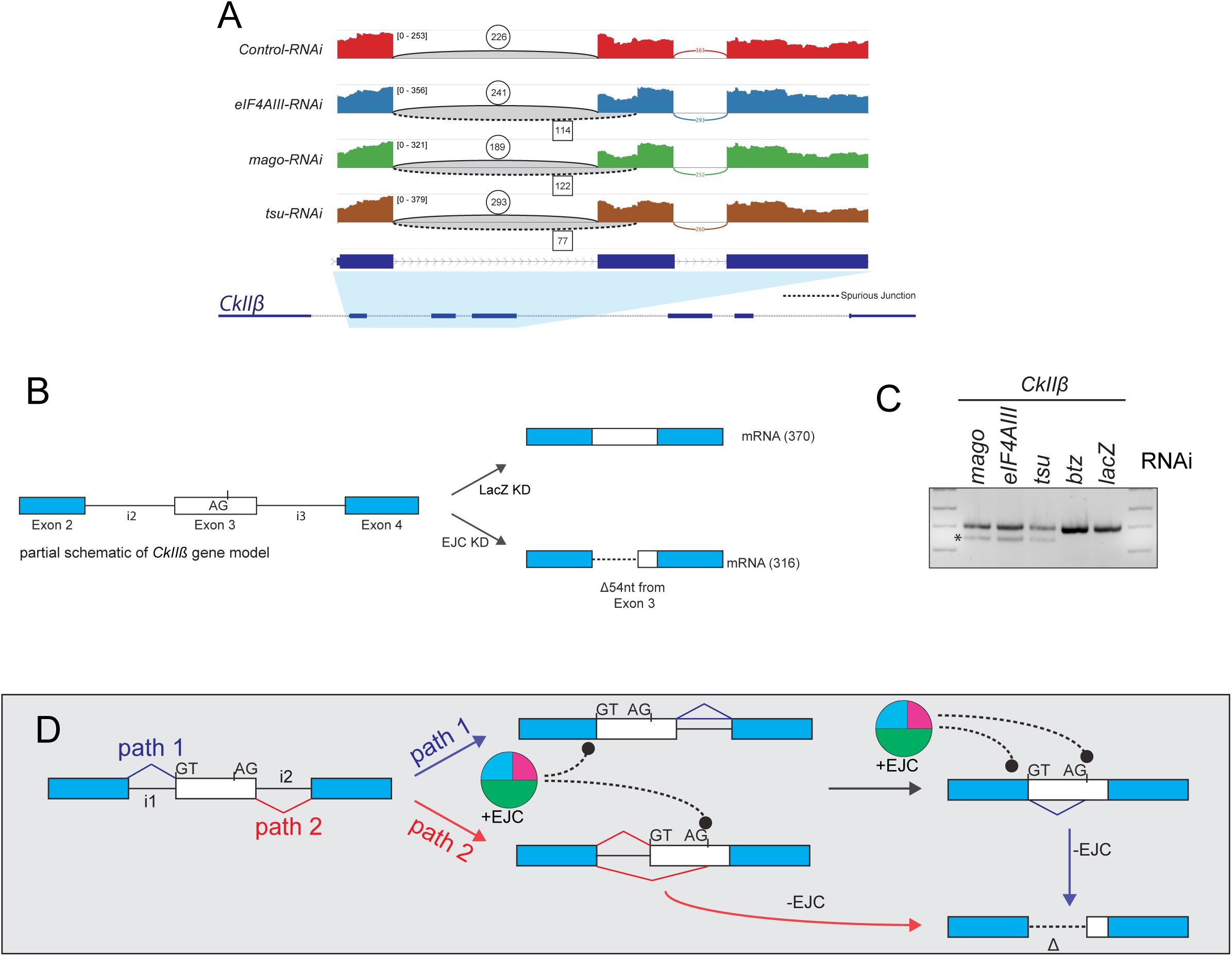
*de novo* splicing on *CkIIß* is a result of dual cryptic splice site activation. (A) Sashimi plot depicting HISAT2-mapped sequencing coverage along a portion of *CkIIß*, which has a cryptic 3’ SS that is activated under core-EJC LOF. Junction spanning read counts mapping to the canonical junction are circled, whereas cryptic junction read counts are squared. Note that spliced reads mapping to the cryptic junction are found in *eIF4AIII, mago* and *tsu* but not the control comparison. (B) Schematic of a *de novo* splicing event detected on the *CkIIß* transcript. (C) Validation of *CkIIß* cryptic 3’ SS activation in core-EJC, but not *btz* or *lacZ* KD conditions (D) Models that explain the *CkIIß* splicing defects. Path 1 and 2 reflect alternate orders of intron removal. Crucially, path 1 leads to EJC-suppressed cryptic splicing on mRNAs using the indicated 5’ recursive splice site and a cryptic 3’SS, whereas path 2 can also produce a splice defect after removal of intron 2.

**Figure 4 - figure supplement 2.**
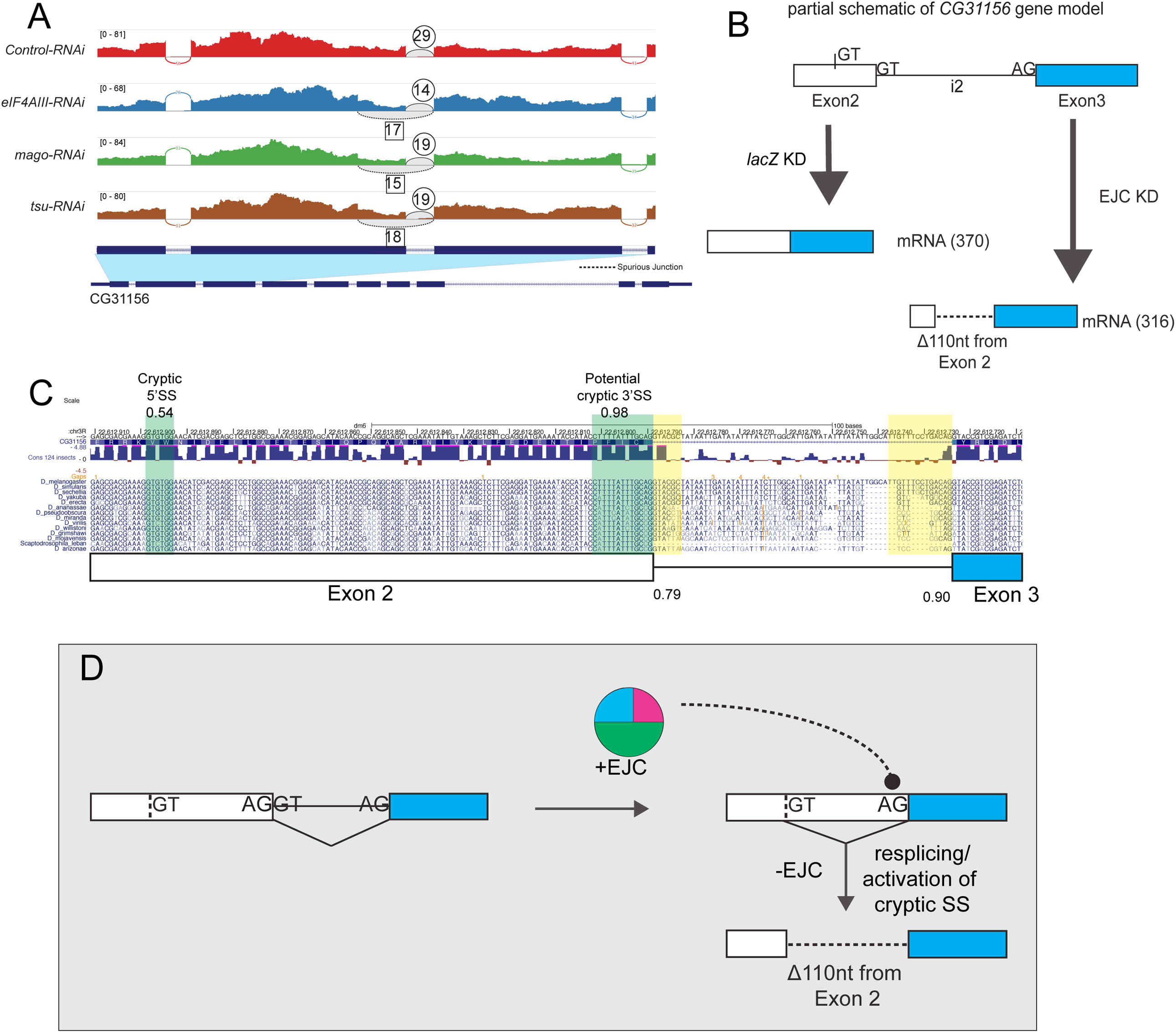
*de novo* splicing on *CG31156* is a result of dual cryptic splice site activation. (A) Sashimi plot depicting HISAT2-mapped sequencing coverage along a portion of *CG31156*, which has a cryptic 5’ SS that is activated under core-EJC LOF. Junction spanning read counts mapping to the canonical junction are circled, whereas cryptic junction read counts are squared. Note that spliced reads mapping to the cryptic junction are found in *eIF4AIII, mago* and *tsu* but not the control comparison. (B) Schematic of a *de novo* splicing event detected on the *CG31156* transcript. (C) Conservation of the cryptic 5’ SS (NNSPLICE score of 0.54) and a potential 3’ recursive splice site (NNSPLICE score of 0.98) found on the *CG31156* transcript highlighted in green, relative to the gene model. Conservation of the splice site is depicted using the multiple alignment format on the UCSC genome browser, as well as phyloP scores. Canonical splice sites are highlighted in yellow. (D) Model of activation of dual cryptic splice sites on the *CG31156* transcript. Activation of the cryptic 5’ SS with an additional cryptic 3’ recursive splice sites leads to deletion of 110 nt of mRNA.

**Figure 4 - figure supplement 3.**
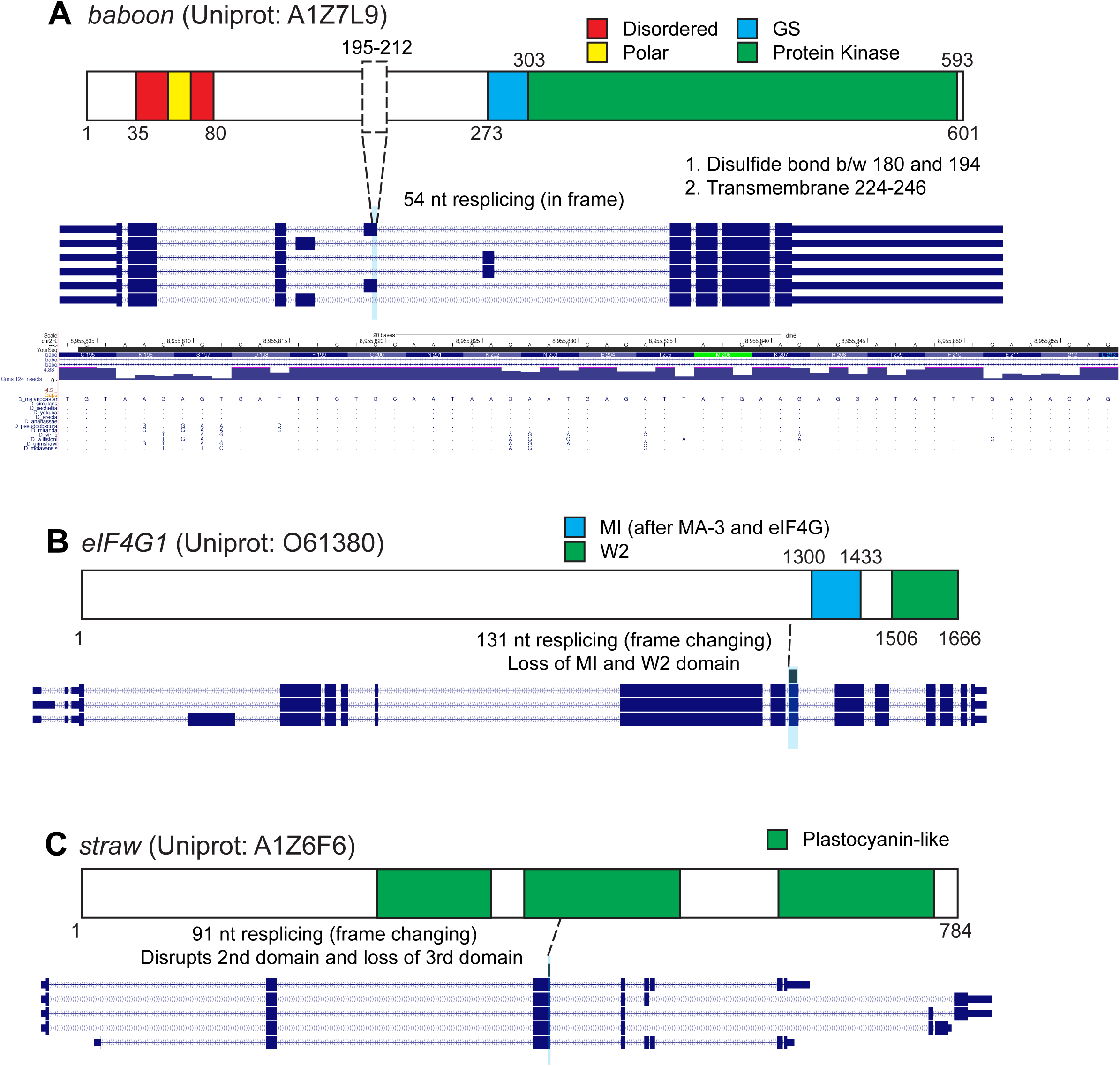
Resplicing on mRNAs alters translated proteins. (A-C) Protein and transcript structures are schematized and the location of cryptic resplicing highlighted in blue. (A) Resplicing on *baboon* leads to a 54 nt deletion of the mRNA and an 18 amino acid deletion. The deletion does not overlap known domains. Conservation plots for deleted 54 nt region is included. (B) Resplicing on *eIF4G1* leads to a 131 nt deletion, leading to a change in reading frame and truncation of the C terminal domains of *eIF4G1*. Importantly, critical domains required for *eIF4G1* function are lost due to resplicing. (C) Resplicing on *straw* leads to a 91 nt deletion, leading to a change in reading frame and truncation of the protein. Importantly, 2 of 3 Plastocyanin-like domains are lost due to transcript defects.

